# SimMiL: Simulating Microbiome Longitudinal Data

**DOI:** 10.1101/2024.03.18.585571

**Authors:** Nicholas E Weaver, Audrey Hendricks

## Abstract

0.

**Motivation:** The quantity of statistical tools designed for omics data analysis has grown rapidly with the ability to collect large sets of human health data, particularly longitudinal data sets. Most tools are assessed for performance using simulated datasets constructed to mimic a handful of relevant characteristics from real world data sets. Consequently, the simulated data sets, and their respective simulation frameworks, are too narrow in scope to qualify as a standard for assessment in longitudinal omics analyses.

**Results:** Here we present the flexible and accessible simulation framework and software package called SimMiL (**Sim**ulating **Mi**crobiome **L**ongitudinal data) capturing three general components of longitudinal microbiome data: (i) absence/presence of microbes, (ii) individual microbe abundance, and (iii) microbiome community composition over time. The framework is assessed by replicating the Type I error and Power analyses of a broad range of statistical tools (MirKAT, repeated measures permANOVA, and a modified kernel association test).

**Software Avaliability:** The simulation framework is at https://github.com/nweaver111/SimMiL

## 1. Introduction

Microscopic organisms form communities in all types of environments, impacting the ecological systems in which they exist. These communities, called microbiomes, help to regulate health and support immune responses related to inflammatory bowel disease, metabolism, and the production of vitamins^1^ when the environment is the human body. Furthermore, identifying longitudinal relationships between changes in microbiomes with changes in human health provides additional insights into the processes and mechanisms of health and disease. However, finding these longitudinal relationships requires the development of statistical methods, which, in turn, require realistic data simulations for method evaluation and comparison. Reproducible simulation frameworks for generating longitudinal microbiome data do not adequately model realistic microbiome community structures over time, leading to inaccurate method assessments and possibly inaccurate biological conclusions when studying longitudinal hypotheses.

Three important characteristics of longitudinal microbiome communities include: (i) the absence/presence of individual microbes, (ii) the abundance of each microbe, and (iii) the overall compositional structure of the microbiome^2–4^. All three characteristics are evident when studying a matrix of counts representing the microbiome. Here, each row of the matrix represents one microbiome sample, with elements of the vector corresponding to the abundance of a microbe.

These vectors of counts can be created by combining microbes into genetically related groups classified at a taxonomic level (e.g., genus or species), denoted generically as operational taxonomic units (OTU). In a study with *N* microbiomes gathered from *n* human individuals across *T* timepoints, it is common for a handful of OTUs to be present in a subset of microbiomes and to leave and re-appear at varying abundances over time^5^. As abundances of individual OTUs change, the structure of the overall microbiome is altered, with some OTUs accounting for larger portions of the entire community than previously seen. Thus, the study of how these compositional structures change and adapt over time is essential for understanding relationships with health. It is reasonable to utilize ecological measures of diversity directly within a simulation framework to simulate specific microbiome structures. Two commonly used diversity measures are: (a) diversity within a single microbiome (*alpha diversity*), and (b) diversity between pairs of microbiomes (*beta diversity*)^6^. Both definitions are general, allowing for the use of different metrics depending upon desired interpretation.

Currently, two simulation frameworks dominate the field when generating longitudinal microbiome datasets from real data: (1) simulating individual OTU counts to create initial time point values that are perturbed with temporal trends and (2) simulating compositional microbiome data with compositional vectors and then perturbing these vectors with temporal trends. Framework (1) directly models characteristics (i) and (ii) of microbiome data and is often employed through zero-inflated mixture models when generating data for a single OTU^2,7–10^. Longitudinal trends are then simulated for OTU abundance by modifying parameters in the mixture model at each timepoint. The R package ZIBR^2^ uses a zero-inflated beta regression model to simulate individual OTUs following this framework. However, simulation framework (1) is limited to the simulation and analysis of a single OTU and cannot be used to accurately study compositional microbiome structures (characteristic (iii)).

In contrast, Framework (2) provides an option for data simulation that includes microbiome structure but does not directly simulate the number of zeros in the data (characteristic (i)). Most commonly, the Dirichlet-multinomial distribution is used for framework (2)^3,4,11,12^. In this framework, the Dirichlet component models community structure at an initial timepoint by simulating compositional vectors (i.e., vectors that sum to 1) for individuals. In this setting, the compositional vectors serve as a measure of alpha diversity for the microbiome. These compositional vectors are then used in the multinomial component to simulate counts representing OTU abundances. Future timepoints are simulated by randomly altering (which we will call perturbing) the OTU abundances between timepoints.

Currently, no longitudinal microbiome simulation software or reproducible framework captures all three components of realistic microbiome data and instead either captures presence/absence (characteristic (i); Framework 1) or community structure (characteristic (ii); Framework 2). Here we present the flexible and accessible simulation framework and software package called SimMiL (**Sim**ulating **Mi**crobiome **L**ongitudinal data). SimMiL enables easy modification of simulation parameters to create a wide breadth of realistic longitudinal microbiome datasets. In section 2, the simulation framework is presented, followed by examples of simulated data sets generated from the framework in section 3. Section 4 consists of a discussion about SimMiL along with remaining gaps within simulation frameworks for longitudinal microbiome data.

## 2. Simulation Framework (1,588 words)

SimMil simulates a zero-inflated count matrix ***X***^**\***^ with dimensions *nT* × *M*, where *n* is the number of individuals, *T* is the number of timepoints, and *M* is the number of OTUs being simulated. The framework simulates at the observational level, generating the baseline count vector, ***x***_*i*1_, for individual *i* at timepoint 1 using the Dirichlet-Multinomial distribution. User-defined longitudinal trends for the relative abundance of OTUs are used to construct initial count vectors 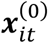 for individual *i* at timepoints 1 < *t* ≤ *T* by using 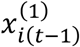 . Finally, a zero-adjustment algorithm is applied to construct the final count vectors 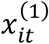 to control the presence and absence of individual OTUs over time. The following subsections outline two important aspects of SimMiL: (2.1) creating temporal trends in an individual and (2.2) the algorithm to control zero-inflation within an observation. The remaining details of the framework are included as supplemental material (Supplemental 1).

### 2.1 Creating Temporal Trends in individual *i*

The baseline count vector for individual *i, x*_*i*1_, is simulated from a Dirichlet-Multinomial distribution as previously done within Framework 2 (See S1 for more details). This vector is converted into a relative abundance vector, ***a***_*i*1_, by dividing each element of *x*_*i*1_by the total number of counts in *x*_*i*1_. A subset of relevant OTUs *S* ⊂ *M* is selected by the user to receive a similar longitudinal trend. This trend is constructed through user-defined weights for timepoints 1 < *t* ≤ *T*, denoted as *w*_*t*_, which are used to convert the initial relative abundance vectors 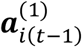 into the vectors 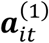 . These relative abundance vectors are then updated through an iterative process to ensure the final relative abundance vector, ***a***_*it*_, maintains the sum-to-one requirement for compositional vectors. An initial count vector for timepoint *t*, 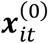, is then simulated from a multinomial distribution with probability vector ***a***_*it*_. The three-step process is described in the following subsections: (2.1.1) Selecting a subset of OTUs *S* ∈ *M*, (2.1.2) defining weights *w*_*t*_, and (2.1.3) maintaining the sum-to-one requirement.

#### 2.1.1 : Selecting a subset of OTUs

Selecting a subset *S* of OTUs depends upon the purpose of the data simulation. For simplicity, the SimMiL framework orders OTUs by average relative abundance across all individuals from the count matrix in the real data set. The simulation parameter λ splits the ordered list of OTUs into rare (low average relative abundance) and common (large average relative abundance) groupings. The simulation parameter *η* controls the proportion of rare or common OTUs that are randomly assigned to the set *S*. Only one subset *S* is selected in SimMiL and it can be one of 4 constructions: (1) only rare OTUs, (2) only common OTUs, (3) a random mixture of rare and common OTUs, or (4) every *d*^*th*^ OTU from the ordered lists.

#### 2.1.2 : Defining the weight for longitudinal changes, w_t_

Known longitudinal trends are incorporated into the elements of ***a***_*it*_ that correspond to the OTUs in *S*. These trends are constructed with weighted changes between time-consecutive relative abundance vectors, as defined in Equation 2.5. The relative abundance for OTU *s* ∈ *S* at timepoint *t* − 1 is multiplied by the time-dependent weight *w*_*t*_. For example, a simulated dataset with *T* = 4 timepoints has three weights: *w*_2_, *w*_3_, and *w*_4_. The relative abundance for OTU *s* ∈ *S* for individual *i* at timepoint 3, denoted as 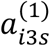, is the product 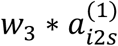. The relative abundance for OTUs *m* ∈ *S*^*c*^ are left unadjusted at this step. Further commentary on selecting values for *w*_*t*_ are found in the discussion section.

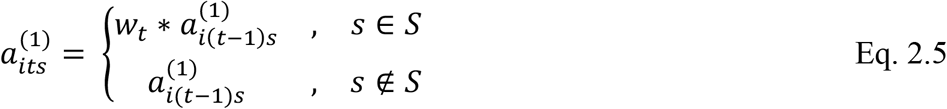

#### 2.1.3 : Maintain sum-to-one constraint

The initial relative abundance vector for individual *i* at timepoint 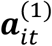, was constructed in section 2.1.2. This vector must be adjusted to ensure the sum-to-one constraint for compositional vectors is valid 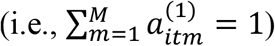. An iterative process that updates the elements of 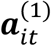 to create the new relative abundance vector 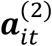 is employed until the vector sum is within a user-defined threshold, *V*, of 1. The algorithm ensures the vector sum gets closer to 1 at each subsequent iteration (see equation 2.6).

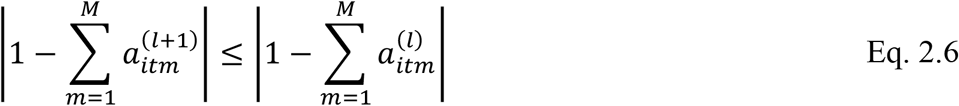

Given an iteration *l* ≥ 1 for the algorithm, the adjustment value 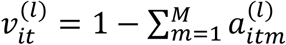 is calculated to determine how far away the current vector sum is from one. Elements of the relative abundance vector 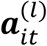 are modified by slightly increasing (or decreasing) values of randomly selected OTUs. The modified vector is stored as 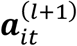 and the adjustment value 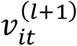 is calculated and compared to the threshold value *V*. The process continues until iteration *L*, the first iteration where 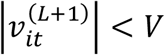. The final relative abundance vector, ***a***, is defined to be equal to the modified vector 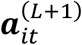. Additional details for modifying the relative abundance vectors and determining *V* are described in detail in supplemental section S.1.2.1.

### 2.2 Controlled zero-inflation within an observation

For timepoints *t* > 1, the initial count vector 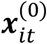 is simulated from a multinomial distribution with mean probability vector ***a***_*it*_. The presence or absence of individual OTUs is then addressed at the observational level starting with individual 1 at timepoint 2. The following process iterates through all *n* individuals before moving to the next time point, eventually ending with subject *n* at timepoint *T*. In general, for individual *i* at time point *t*, 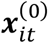 is updated in a three-part process: (1) determine the expected number of absent OTUs in the observation, (2) select a subset of OTUs to be absent in the observation, and (3) redistribute the counts of 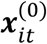 to create the final count vector, 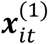.

#### 2.2.1. : Select the number of absent OTUs

A truncated normal distribution is used to simulate ô_*it*_, the desired number of absent OTUs in 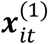. The distribution is defined by a user-selected mean that changes by timepoint, *o* _*t*_,and a variance component that is equal to the sample variance of the number of zero counts across all individuals in the real-world dataset. The distribution is truncated to ensure the number of absent OTUs never exceeds a user-defined feasible region. Additionally, the desired number of absent OTUs, ô_*it*_, is rounded to the nearest whole number to adjust for the continuous nature of the truncated normal distribution.

#### 2.2.2: Select Absent OTUs

A selection algorithm, called the Zero-Adjustment Selection Algorithm, is applied to all observations with timepoint 1 < *t* ≤ *T* to construct the set of absent OTUs within an observation, denoted as γ_*it*_. Because counts are constructed in time-order, OTUs that are absent in *t* − 1 (i.e., γ_*i*(*t*−1)_) are assessed first to incorporate a user-specified level of autocorrelation in OTU absence. The OTUs of γ_*i*(*t*−1)_ are ordered from lowest relative abundance to largest relative abundance with respect to ***a***_*it*_. An OTU remains absent at timepoint *t* with user-defined probability *p*_*m*_, where *p*_*m*_ depends upon the classification of OTU *m* as rare or common (see section 2.1.1). Otherwise, the OTU will be placed in the set of present OTUs, ρ_*it*_. If |γ_*it*_| ≥ ô_*it*_ the algorithm terminates and all unassigned OTUs are placed in ρ_*it*_. However, if |γ_*it*_| < ô_*it*_ the OTUs from ρ _(*i t*−1)_ with a count of 0 in 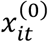 are assigned to γ _*it*_ in order of relative abundance in ***a***_*it*_ (lowest to largest). If at any point |γ_*it*_| = ô_*it*_ the algorithm terminates and remaining OTUs are placed in ρ_*it*_. However, if |γ_*it*_| < ô_*it*_ is still true, the remaining unassigned OTUs are ordered from lowest to largest relative abundance and placed into γ_*it*_ with probability 1 − *p*_*m*_ until |γ_*it*_| = ô_*it*_ or all OTUs have been assigned.

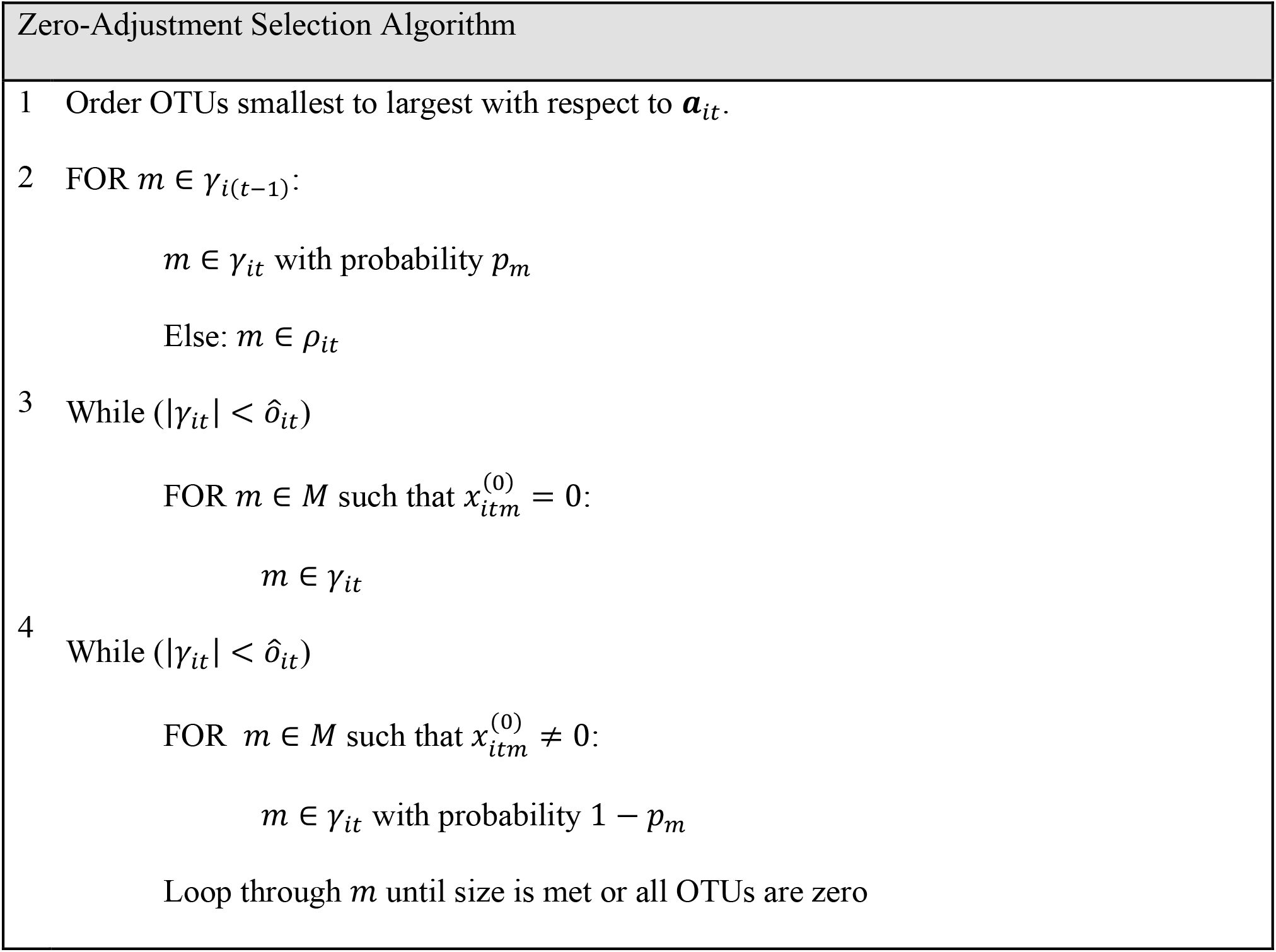

#### 2.2.1. : Redistributing Counts

The constructed sets for absent OTUs, γ_*it*_, and present OTUs, ρ_*it*_, are used to convert the count vector 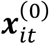 into the final count vector 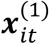 such that the vector sum of 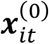 is as close as possible to the vector sum of 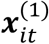. An algorithm for redistribution is created to accomplish the goal. First, the total number of counts, *C*, in 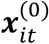 that need reassigned to different OTUs to satisfy γ_*it*_ and ρ_*it*_ is determined. If an OTU was originally simulated to have a non-zero count in 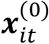, but has been assigned to be absent (i.e., OTU is in γ_*it*_), then *C* is increased by the OTUs count in 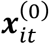. Alternatively, if an OTU was assigned to be present (i.e., OTU is in ρ _*it*_ ) but has an initial count of zero in 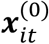, it is given a count of 1 and *C* is decreased by 1. If *C* is a positive number after all OTUs have been adjusted, the remaining counts are re-assigned to OTUs in ρ_*it*_ by using a multinomial distribution with total number of counts equal to *C* and probability vector equal to the relative abundances in ***a***_*it*_ that correspond to the OTUs in ρ_*it*_. The process is summarized in the Algorithm for Redistribution figure.

##### Algorithm for Redistribution

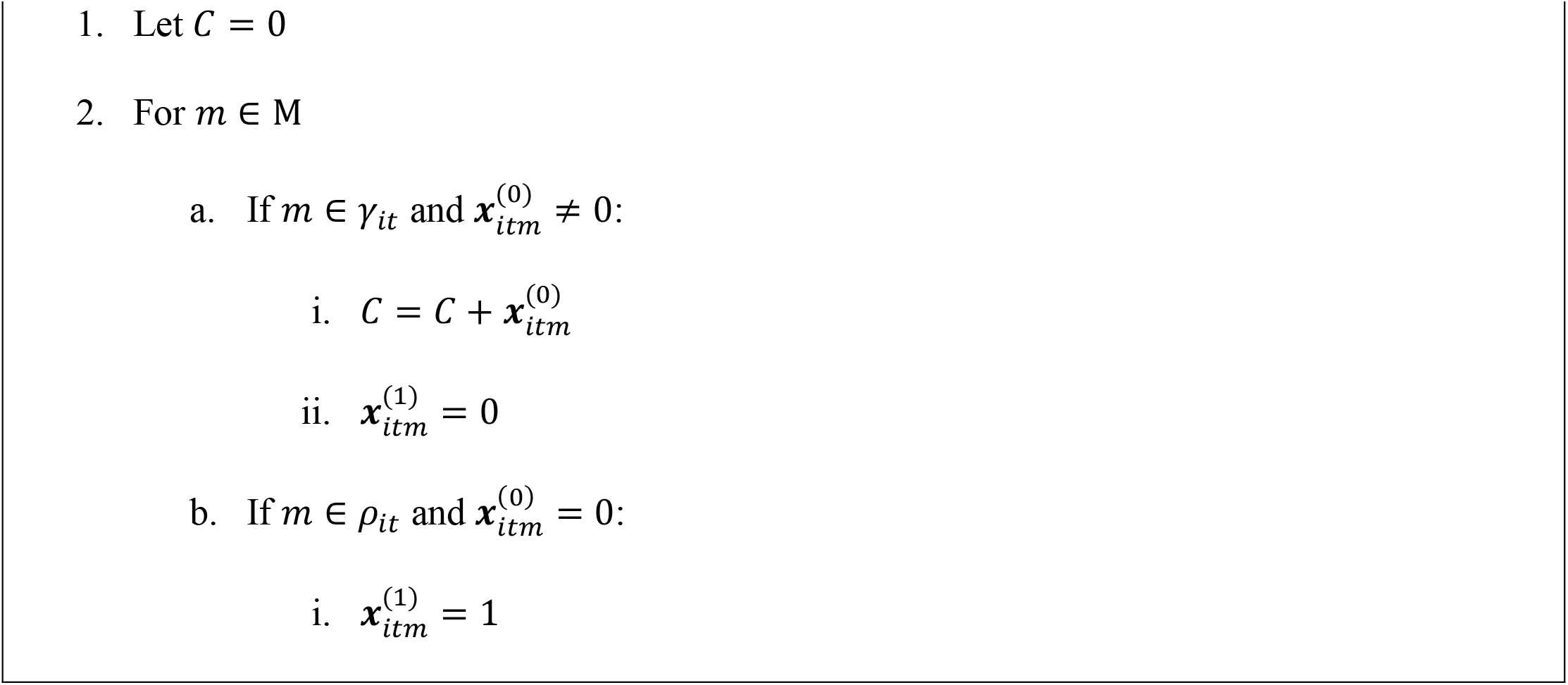
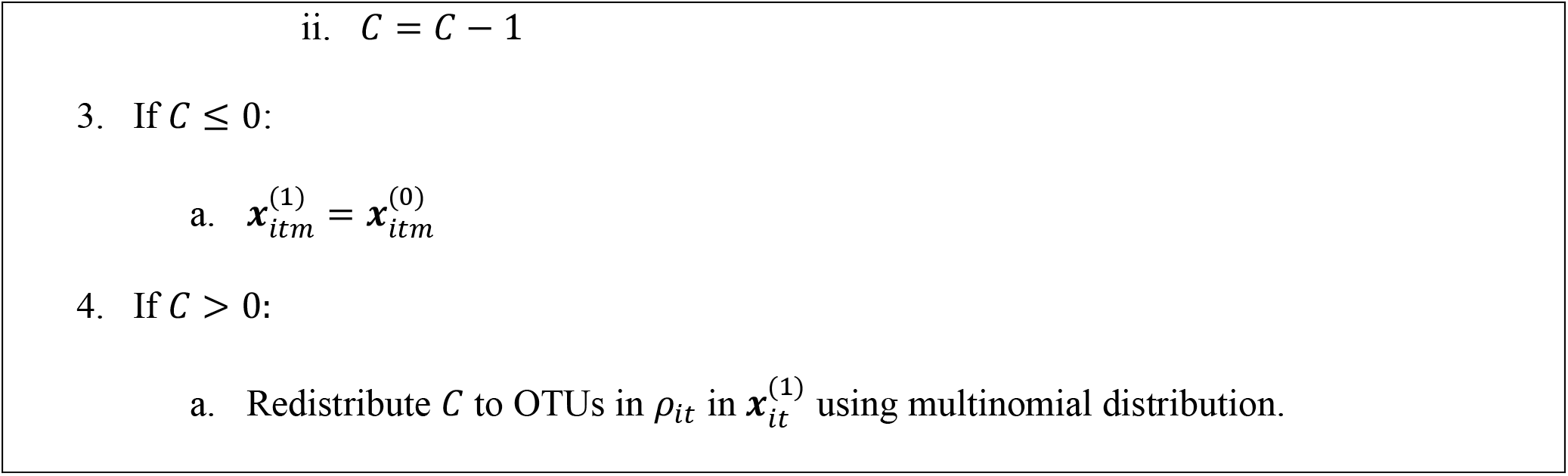

The Algorithm for Redistribution completes the construction of the simulated observation, 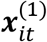. Completing the process across all observations produces the final simulated dataset; a matrix of OTU counts, *X*^*^. The collection of simulated observations contains a known longitudinal trend within a subset of OTU counts as well as a known trend in the average number of absent OTUs over time. Additionally, the simulated data maintains important properties of compositional vectors that mimics the community structure of real microbiome data.

## 3. Results

The SimMiL framework, with corresponding R package (S.2), offers flexibility when simulating longitudinal microbiome data. The framework contains many parameter combinations, all of which would be unreasonable to explore in this manuscript alone. Instead, simulation results are demonstrated for three important parameters: the time change vector (*w*_*t*_), the zero trend vector (*o*_*t*_), and the subset of OTUs with known temporal trend (*S*). Each parameter is considered at multiple values and scenarios as described in the following two subsections. In each scenario, a real-world gut microbiome dataset consisting of 118 OTUs (species level) from pregnant women in Guatemala (CITE: Women First) is used to simulate *n* = 100 individuals across *T* = 5 timepoints. The relative abundance cutoff for rare and common OTUs is set to λ = 0.01, the proportion of rare/common OTUs selected for *S* is set to *η* = 0.5, the probability a rare taxa changes from absent to present between consecutive timepoints is *p*_*R*_ = 0.2 while the probability a common taxa changes from absent to present between consecutive timepoints is *p*_*C*_ = 0.7.

### 3.1 Longitudinal trends through weight parameter *w*_*t*_ and subset *S*

Three types of longitudinal trends are considered: monotonic increasing, monotonic decreasing, and non-monotonic. The monotonic increasing trend doubles the relative abundance of OTUs in *S* with a weight vector of 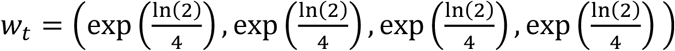. A similar formulation is used to create the monotonic decreasing trend, however the relative abundance is cut in half with 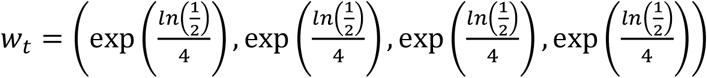. Finally, the non-monotonic dataset forces the relative abundance of OTUs in *S* to double before returning to the initial relative abundance value of *t* = 1 with weight vector 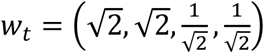.

To incorporate potential differences due to the selection of *S*, two data sets are simulated for each trend based upon the construction of the subset: half the common OTUs, or 30 OTUs (about ¼ of all OTUs) with varying levels of relative abundance. The trend in absent OTUs (*o*_*t*_) for each dataset is set to be constant at all timepoints such that the mean number of absent OTUs matches the mean number of absent OTUs in the real-world dataset (default value in SimMiL). The three longitudinal trends are assessed using visualization tools in the SimMiL package to understand the impact of *w*_*t*_ and *S* on the average microbiome composition over time.

Figure 3.1.1 shows the average relative abundance of perturbation groups (either *S* or *S*^*c*^) across all individuals at each timepoint using stacked bar graphs. The three longitudinal trends for *w*_*t*_ are split by row and are most evident in the simulated data when the subset *S* is comprised of only common OTUs (left column of Figure 3.1.1). In contrast, the temporal trends are less pronounced in the stacked bar graphs when a mixture of rare and common OTUs are selected for *S*, implying a smaller component of the total microbiome is changing at each timepoint.

**Figure 3.1.1.**
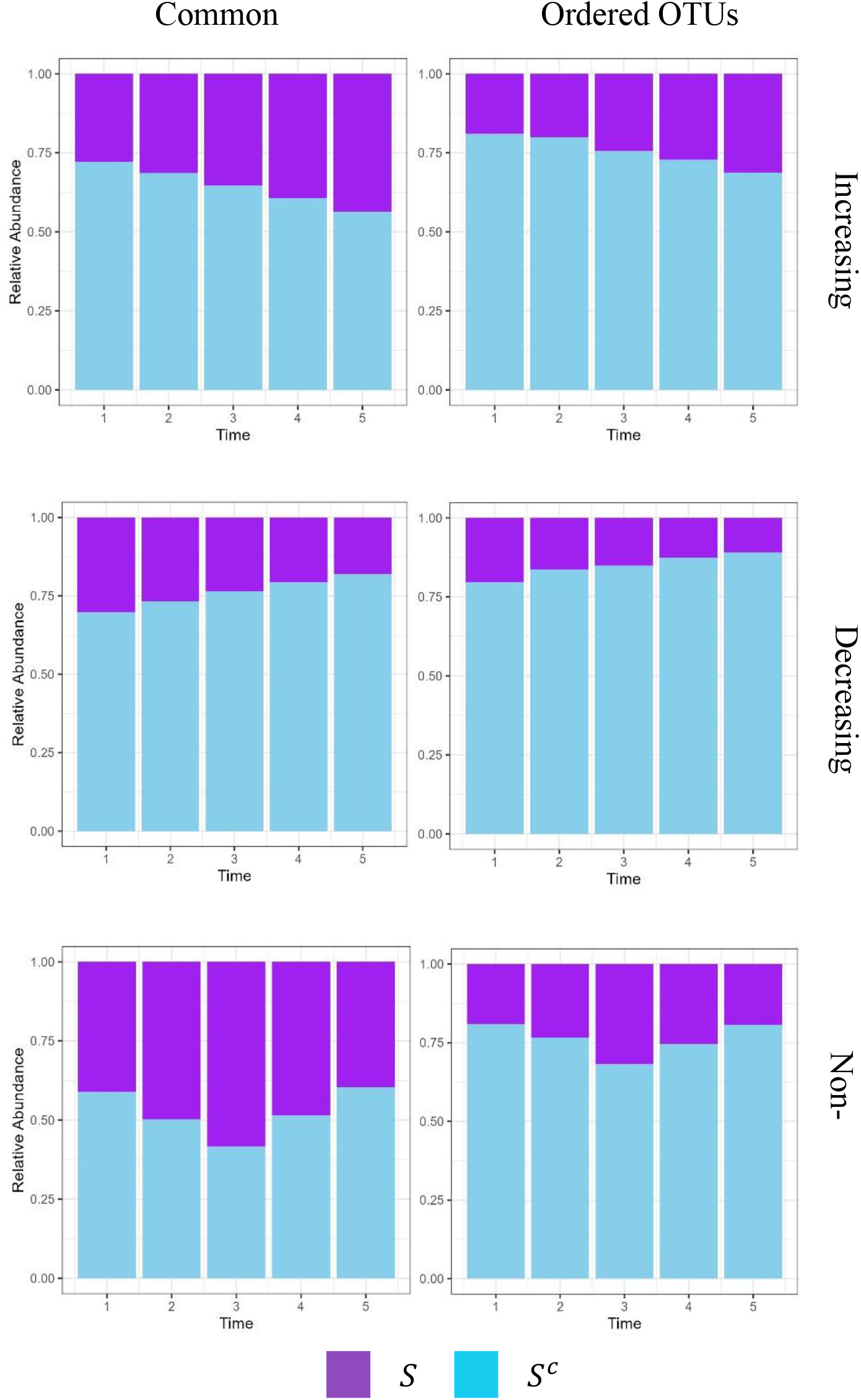
Longitudinal Trends with Constant Zeros by Perturbation Grouping. Stacked bar graphs representing the average total relative abundance of OTUs in *S* and *S*^*c*^ across time. The left column represents scenarios where *S* is half of the common OTUs while the right columns represents scenarios where *S* is 30 OTUs of varying relative abundance. The top row is all scenarios with monotonic increasing trend, the middle row has monotonic decreasing trend, and the bottom row has a non-monotonic trend.

Figure 3.1.2 shows the average relative abundance of each OTU (118 segments) across all individuals at each timepoint using stacked bar graphs. These plots highlight the temporal behavior of relative abundance in the most common OTUs. For scenarios where only common OTUs are in *S* (left column of Figure 3.1.2), there is a clear trend over time that matches the simulation parameter *w*_*t*_ (monotonic increasing, monotonic decreasing, or non-monotonic). In contrast, the scenarios where *S* contains both rare and common OTUs (right column of Figure 3.1.2) show the impact of randomly redistributing expected relative abundance values to maintain the sum-to-one requirement. Most notably, when common OTUs are not in *S*, it is likely to see random fluctuations in common OTUs between timepoints because they are only used for redistribution in SimMiL.

**Figure 3.1.2.**
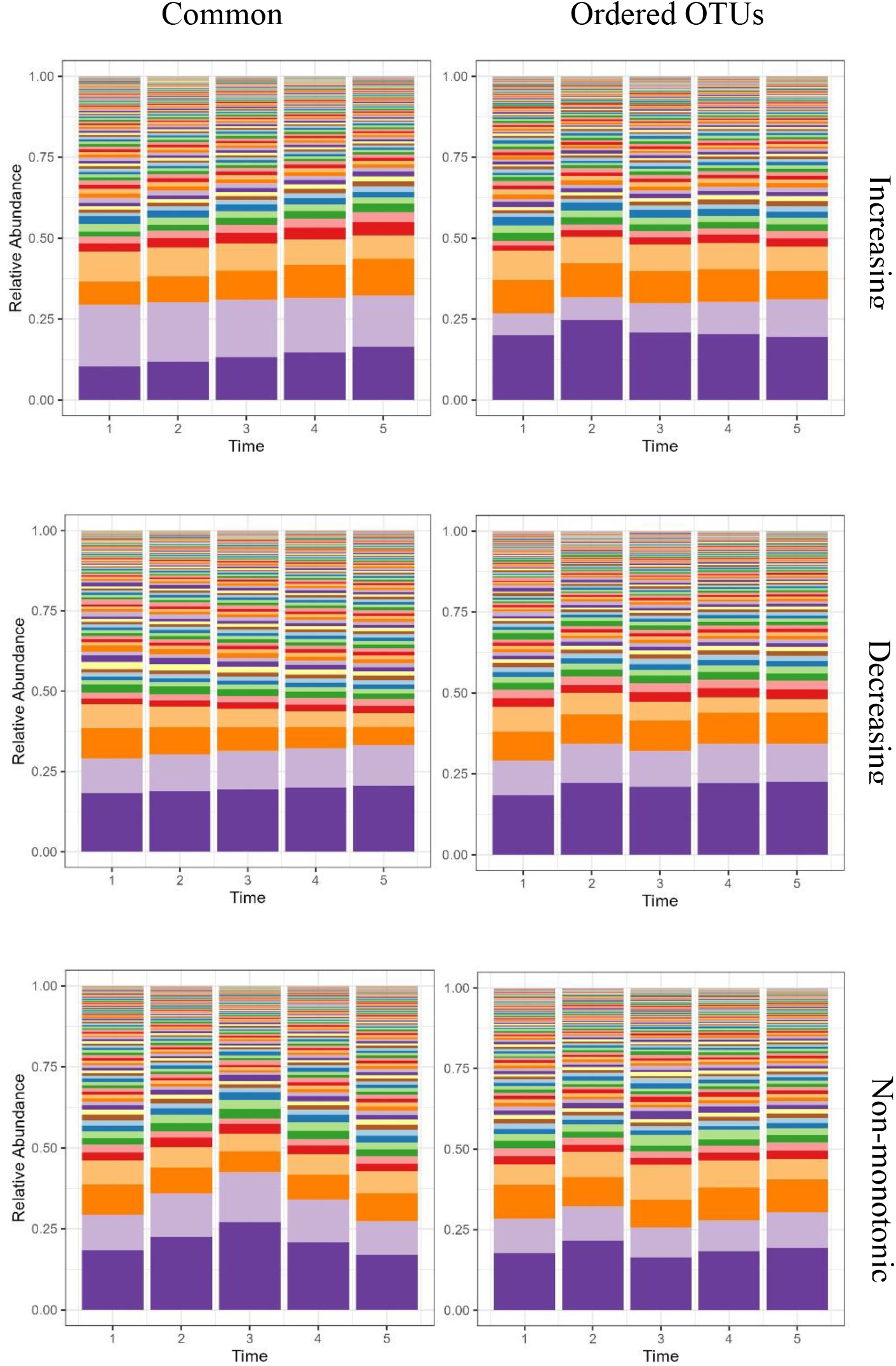
Longitudinal Trends with Constant Zeros by Taxa. Stacked bar graphs representing the average relative abundance of OTUs across time. The left column represents scenarios where *S* is half of the common OTUs while the right column represents scenarios where *S* is 30 OTUs of varying relative abundance. The top row is all scenarios with monotonic increasing trend, the middle row has monotonic decreasing trend, and the bottom row has a non-monotonic trend.

Figure 3.1.3 shows the average beta diversity value between observations from the same individual in all 6 scenarios. Because SimMiL simulates an observation by using the previous timepoint, an autocorrelation structure is expected between microbiome structures. However, when comparing the pairwise similarities of all 5 timepoints within an individual, there do not appear to be any longitudinal trends. This is true regardless of the simulation scenario, s seen in Figure 3.1.3.

**Figure 3.1.3.**
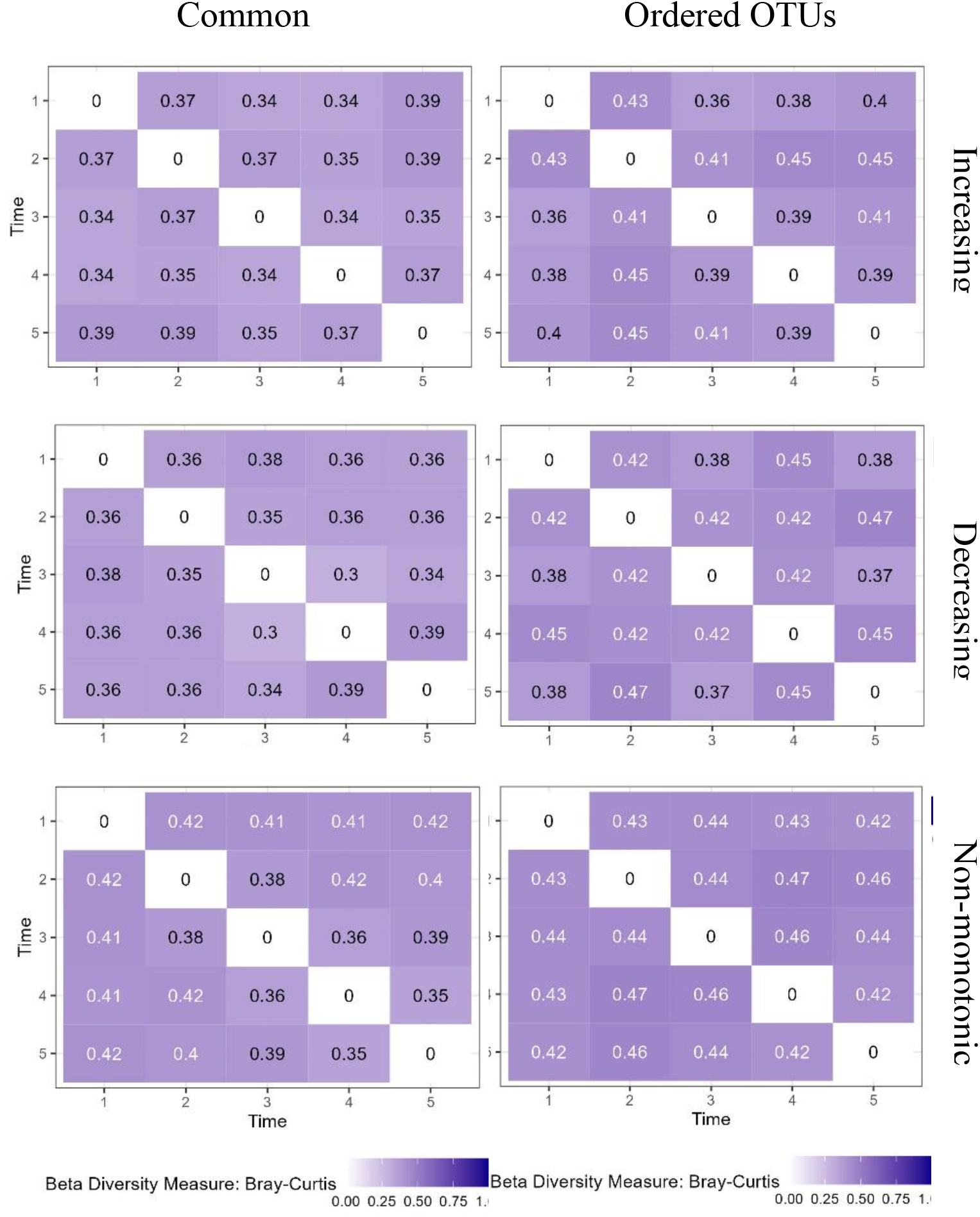
Longitudinal Trends with Constant Zeros by Beta Diversity. Heatmaps representing the average beta diversity between observations within an individual, across all individuals. The left column represents scenarios where *S* is half of the common OTUs while the right column represents scenarios where *S* is 30 OTUs of varying relative abundance. The top row is all scenarios with monotonic increasing trend, the middle row has monotonic decreasing trend, and the

### 3.2 Altering the structure of absent OTUs

Three types of trends in zero counts (*o*_*t*_) are considered for comparison: monotonic increasing, monotonic decreasing, and non-monotonic. The monotonic increasing trend increases the mean number of absent OTUs by 5 at each timepoint, 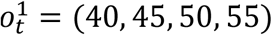, while the monotonic decreasing trend decreases the number of absent OTUs by 5 each timepoint, 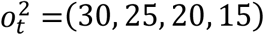. Finally, the non-monotonic dataset forces the mean number of absent OTUs at each timepoint to reach half of all OTUs at timepoint *t* = 3 before returning to the mean number of absent OTUs from timepoint *t* = 1, 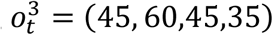. The mean number of absent OTUs at *t* = 1 is found by using the mean number of absent OTUs in the real-world data set.

To focus solely on the impact of zero counts, the temporal trend vector is set to be constant across timepoints, *w*_*t*_ = (1,1,1,1). In other words, the relative abundance of each OTU is not adjusted between timepoints. Consequently, the selection of *S* is not meaningful in this setting and is set to the SimMiL default.

**Figure 3.2.1** shows boxplots binned by relative abundance thresholds. Within a bin is one boxplot for each simulated timepoint. The boxplots represent the proportion of OTUs within an observation with relative abundance matching the bin threshold. The three simulation structures (increasing, decreasing, non-monotonic) are represented as rows in the figure. Because these scenarios differ by absent OTU trends, the most informative bin of each panel is that of zero (left-most bin in Figure 3.2.1). As time increases within the zero bin, the three simulated trends are explicitly seen by vertical shifts of the boxplots. It is also evident in Figure 3.2.1 that any temporal trends in the zero bin are mirrored by inverted temporal trends in the next smallest bin, when the relative abundance is between 0 and 0.001. This is consistent across all three types of trends.

**Figure 3.1.2.**
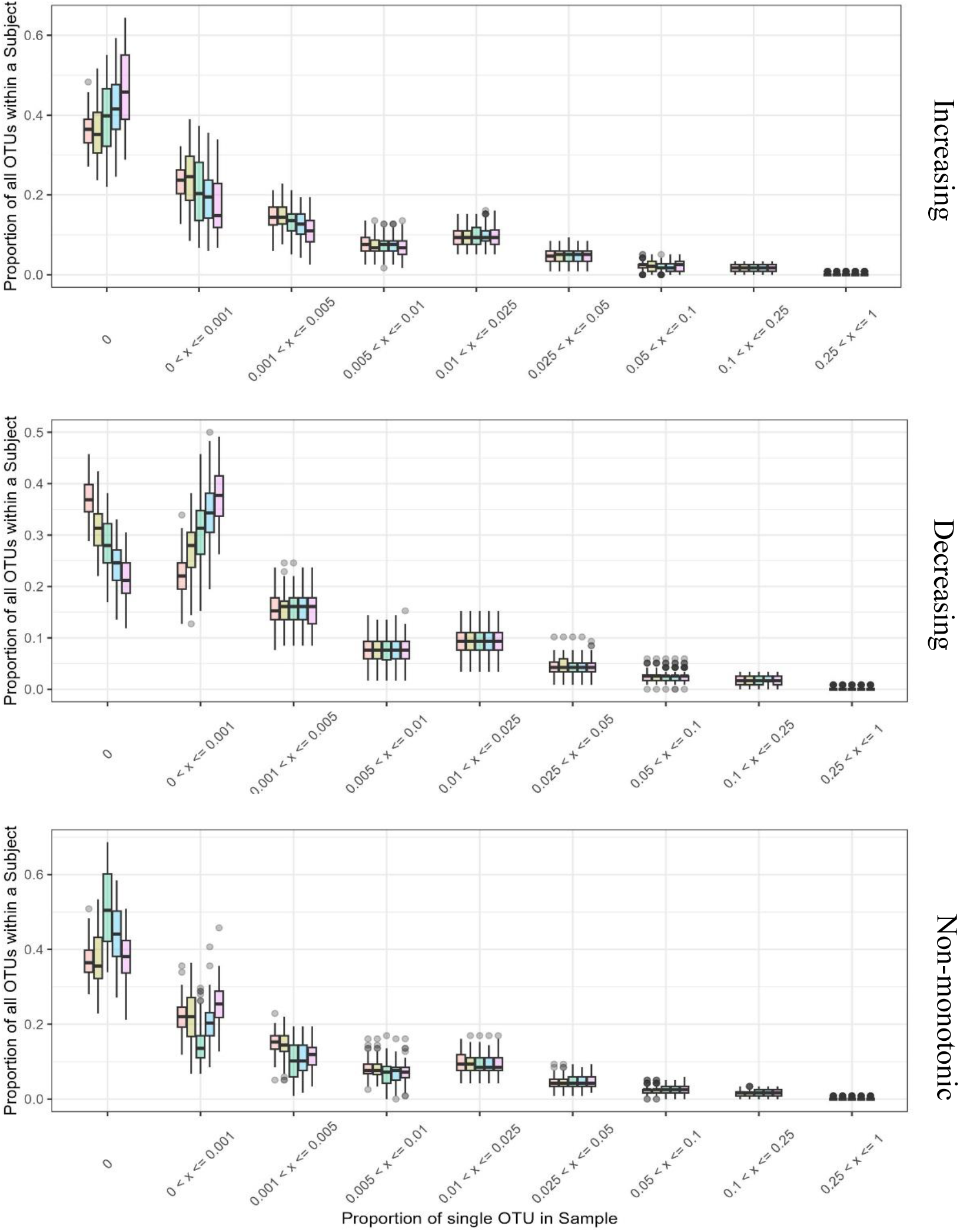
Longitudinal Trends in Absent OTUs. Binned boxplots representing the proportion of all OTUs (vertical axis) within an observation with relative abundance at different levels of rare and common (horizontal axis) split by timepoint. The top row is the scenario with monotonic increasing trend in absent OTUs, the middle row has monotonic decreasing trend in absent OTUs, and the bottom row has a non-monotonic trend in absent OTUs.

## 4. Discussion

SimMiL extends the popular Dirichlet-multinomial framework for generating compositional count data at a single timepoint to multiple timepoints. Three important attributes of longitudinal count data are prioritized in the SimMiL framework: (1) easily generating known longitudinal trends within each feature, (2) allowing counts of a feature to switch between zero and non-zero over time, and (3) maintaining the compositional nature of the count data. To our knowledge, no readily available simulation framework and R package for longitudinal microbiome data accomplishes all three aspects. Overall, the SimMiL framework produces biologically appropriate simulated datasets with multiple input parameters that are used to create different types of longitudinal trends in microbiome count data.

Longitudinal trends in SimMiL are constructed through time-dependent weights. These weights must be positive numbers that support a realistic and desired simulated outcome. A weight less than 1 decreases relative abundance between consecutive timepoints whereas a weight greater than 1 increases relative abundance between consecutive timepoints. In most settings, a scenario where the weight is constant across time is appropriate (i.e., *w*_*z*_ = *w*_*y*_ for all *z, y* ∈ {2, …, *T*}), as was shown in section 3. It can be shown that scenarios with constant weight produce OTUs with relative abundances that follow exponential growth (if weight greater than 1) or decay (if weight less than 1) over time. Thus, a simulated dataset where the relative abundance changes by a factor of *W* between timepoints 1 and 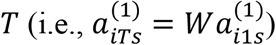 is constructed by setting 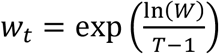 for all *t* ∈ {2,3, …, *T*}. Alternative types of growth, decay, or hybrid change can be studied by assigning different weights at each time point. The flexibility of how weights are selected provides opportunity for almost any reasonable scenario to be simulated through the SimMiL framework.

The behavior of OTUs that are absent is well controlled in the SimMiL framework. The mean and variance for the number of absent OTUs at each timepoint is easily specified through framework parameters. Additionally, parameters that determine the chance an OTU becomes non-zero at a future timepoint provide extra control over the absence and presence behavior of individual OTUs. It is important to discuss that SimMiL controls the absence structure of simulated data by modifying the structure of extremely rare OTUs, as seen in the results of Section 3.2. This behavior is not expected for all microbiome datasets and is a current weakness of SimMiL. Future work to create a new Zero-Adjustment algorithm to better mimic what is seen in real world data is needed.

The role of ecological diversity is another important component when simulating longitudinal microbiome data. Frameworks such as SimMiL and the popular Dirichlet-Multinomial framework utilize compositional vectors to replicate and control alpha diversity within simulated data. Currently, no commonly used simulation framework that simulates count matrices can be used to directly simulate beta diversity values. Although the SimMiL framework is capable of simulating data with realistic beta diversity values as seen in Section 3, it is not clear how to simulate longitudinal trends into these values. Instead, one could experiment with different parameter values until simulating the desired longitudinal trends in beta diversity. A future extension of the SimMiL package, and underlying SimMiL framework, is needed to explicitly simulate count data with known longitudinal trends in beta diversity.

Which OTUs have meaningful and known change over time is foundational to studying relationships between microbiomes and health. Most real microbiomes typically consist of multiple longitudinal trends across several different groupings of OTUs, not just a single trend. Additionally, most subsets of OTUs that have similar longitudinal trends in real datasets also have high phylogenetic relatedness with one another. SimMiL simulates a single longitudinal trend in a subset of OTUs grouped by relative abundance values. Although the SimMiL R package currently offers four options for selecting *S*, it is not currently possible to choose multiple subsets or to pick a subset through relatedness of OTUs. However, because microbiome datasets are constructed by categorizing OTUs with phylogenetic trees, it would be reasonable to select OTUs for subset *S* through a clustering algorithm using phylogenetic relatedness as the distance measure (e.g., the partition around medoids algorithm)^4,12,13^. Future extensions to the SimMiL framework and R package should include input parameters that allow users to pick a list of OTUs for *S*, or to input a phylogenetic tree to form *S*. Likewise, future extensions should provide a way for multiple longitudinal trends to be simulated into the same dataset.

Another important aspect of microbiome data is the total number of OTUs being studied (*M*). One current limitation of the SimMiL framework and R simulation package is the number of OTUs that can be simulated. Because the framework focuses on compositional vectors from real-world data, the number of OTUs generated must match the number of OTUs in that real-world dataset, making it impossible to construct scenarios with different numbers of OTUs. An extension to the framework that allows the user to identify the number of OTUs to generate (either more or less than seen in the real-world data) would be helpful when generating even larger datasets. This could potentially be done by removing the real data from the framework and instead using a parameter input for the total number of OTUs and the proportion of those OTUs that should be rare or common.

Finally, the simulation package is computationally expensive compared to ZIBR and other simulation packages that currently exist. As the number of simulated individuals, or timepoints, increases, the computational time of SimMiL increases linearly. This expense is primarily caused by SimMiL simulating data at the observational level as shown in Section 3.3. However, other factors like inefficient code and the need to calculate the maximum likelihood values for parameter estimates from the real-world data can be solved through future updates to the SimMiL R package. In this way, larger simulation projects can be completed in less time, increasing the utility of the SimMiL framework and R package.

## Supporting information

Supplemental Sections

