## Supplemental Sections for "SimMiL: Simulating Microbiome Longitudinal Data"

Supplemental Materials

### S.1 Methods

#### S.1.1 Simulating the first time point in $\boldsymbol{X}^{(1)}$

The Dirchlet-Multinomial (DM) simulation framework simulates count vectors for individuals at a single time point. Given individual $i$, we simulate the vector of OTU counts $\boldsymbol{x}_{i1}^{(1)}$ at baseline (or timepoint 1) for the initial iteration of SimMiL using the approach described by Zhao et al. The steps of the process include: (1) estimating Dirichlet hyperparameters from a real dataset at a single timepoint ($\boldsymbol{\alpha}$), (2) simulating individual-level baseline OTU proportions, $\boldsymbol{a}_{i1}$, using the Dirichlet distribution and hyperparameters, and (3) generating individual-level baseline OTU counts, $\boldsymbol{x}_{i1}^{(1)}$ using a multinomial distribution with probability parameter equal to $\boldsymbol{a}_{i1}$. In this way, $n$ vectors of baseline counts for $M$ conditionally related OTUs are generated and used to simulate count vectors for subsequent timepoints.

##### *S.1.1.1: Estimating Dirichlet hyperparameters*

Hyperparameters of a Dirichlet distribution are estimated using the DM likelihood function shown in Equation 2.1. The parameters for overdispersion, $\theta$, and the mean compositional vector, $\boldsymbol{\pi}$ (length $M$), are given as the function of a real OTU count matrix, $\boldsymbol{X}$ ($n' \times M$), and corresponding total count vector for each community, $\boldsymbol{\eta}$ ($n' \times 1$). Here the notations $\boldsymbol{X}$, $n'$, and $M$ are used to denote the count matrix, number of observations, and number of OTUs, respectively, from a real-world dataset. To estimate the DM hyperparameters $\theta$ and $\boldsymbol{\pi}$, Equation 2.1 is maximized to find the MLE estimates for both parameters.

$$L(\theta, \boldsymbol{\pi} | \mathbf{X}, \boldsymbol{\eta}) = \prod_{i=1}^{n} \binom{\eta_i}{\boldsymbol{x_i}} \frac{\Pi_{j=1}^{M} \Pi_{r=1}^{x_{ij}} \{\pi_j (1-\theta) + (r-1)\theta\}}{\Pi_{r=1}^{\eta_i} \{1 - \theta + (r-1)\theta\}} \qquad \text{Eq. 2.1}$$

#### *S.1.1.2: Creating Vectors of Proportions by Combining Hyperparameters*

An average relative abundance vector for the real-world data is constructed by combining the MLE estimates $\hat{\boldsymbol{\pi}}$ and $\hat{\theta}$. We denote this new vector as $\boldsymbol{\alpha}$, where the $j^{th}$ element of $\boldsymbol{\alpha}$ is $\alpha_j = \hat{\pi}_j * \frac{(1-\hat{\theta})}{\hat{\theta}}$ [14]. A Dirichlet distribution (Equation 2.2) with parameter $\boldsymbol{\alpha}$ is used to simulate a relative abundance vector for each individual, creating baseline community structures for all individuals.

$$\mathrm{p}(\boldsymbol{a}|\boldsymbol{\alpha}) = \frac{\Gamma(\sum_{m=1}^{M} \alpha_m)}{\prod_{m=1}^{M} \Gamma(\alpha_m)} * \prod_{m=1}^{M} a_m^{\alpha_m - 1} \qquad \text{Eq. 2.2}$$

$$\text{s.t. } \sum_{m=1}^{M} a_m = 1;\ a_m \in (0,1)$$

#### *S.1.1.3: Simulate individual-level baseline OTU counts*

A baseline count vector for individual $i$, denoted as $\boldsymbol{x}_{i1}$, is generated from the multinomial distribution (Equation 2.3) using $a_i$ for the proportion parameter. To simulate differences in read depth between observations, the number of counts to distribute for individual $i$ at time point 1 (aka baseline) is denoted by $\eta_{i1}$, and is simulated from a negative binomial distribution (Equation 2.4). The mean, $\mu$, and standard deviation, $\phi$, parameters for the negative binomial distribution are the mean and standard deviation of total reads across all observations in the real-world dataset.

$$p(\boldsymbol{x}_{i1}|\eta_{i1}, \boldsymbol{a}_{i1}) = \frac{\eta_{i1}!}{\prod_{m=1}^{M} x_m!} \prod_{m=1}^{M} a_{i1m} \qquad \text{Eq. 2.3}$$

$$p(\eta_{i1}|\mu, \phi) = \binom{\eta_i + \phi - 1}{\eta_i} \left(\frac{\mu}{\mu + \phi}\right)^{\eta_i} \left(\frac{\phi}{\mu + \phi}\right)^{\phi} \qquad \text{Eq. 2.4}$$

### S.1.2 Simulating temporal trends

#### *S.1.2.1: Sum-to-One Algorithm*

Iteration $l$ constructs the relative abundance vector $a_{it}^{(l)}$ by assigning segments of $v_{it}^{(l)}$ randomly to the elements of $a_{it}^{(l-1)}$. Initially, the parameters for a multinomial distribution are established by converting $v_{it}^{(l)}$ into the whole number $\Psi_{it}^{(l)} := \text{round}(10^{c} * v_{it}^{(l)})$, where $c$ represents the number of decimal places in the threshold value $V$ (e.g., if $V = 0.000001$, then $c = 6$). Next, the vector $d_{it}^{(l)}$ is simulated from the multinomial distribution with total counts equal to $\left|\Psi_{it}^{(l)}\right|$ and a probability vector equal to $a_{it}^{(l-1)}$. For each $m \in M$, the proposed change for the relative abundance between iterations $l-1$ and $l$ is defined as $D_{itm}^{(l)} := \text{sign}\left(\Psi_{it}^{(l)}\right) * \frac{\left(d_{itm}^{(l)}\right)}{10^{c}}$, and thus the proposed new relative abundance vector is calculated as $a_{itm}^{(l)} = a_{itm}^{(l-1)} + D_{itm}^{(l)}$. If $a_{itm}^{(l)}$ is no longer in the feasible domain of $(0,1)$, the relative abundance for $m$ is not adjusted (i.e., $a_{itm}^{(l)} = a_{itm}^{(l-1)}$). Next, the stopping criteria is checked. If $\left|1 - \sum_{m=1}^{M} a_{itm}^{(l)}\right| > V$ the process moves to iteration $l+1$, otherwise the process terminates with $a_{itm}^{(l)} = a_{itm}^{(L+1)}$. Finally, the relative abundance vector $\boldsymbol{a}_{it} := \frac{\boldsymbol{a}_{it}^{(L+1)}}{\sum_{m=1}^{M} a_{itm}^{(L+1)}}$ guaranteeing the vector sums-to-one with minimal changes between $\boldsymbol{a}_{it}^{(L+1)}$ and $\boldsymbol{a}_{it}$.

### S.2 SimMiLManual

The R Package, SimMiL, provides an easy-to-implement version of the SimMiL simulation framework. SimOTUCounts is the simulation function in the package, while additional functions are used to create data visualizations for count data. The package is introduced below in two parts: (1) Simulating longitudinal microbiome data and (2) visualizing longitudinal microbiome data.

#### S.2.1 Simulating Longitudinal Microbiome Data.

SimOTUCounts() utilizes a cross-sectional OTU count matrix from a real, user-provided data source to simulate longitudinal microbiome count data. Function inputs control for (1) longitudinal trends, (2) the average number of absent OTUs, and (3) the probability an OTU re-enters the microbiome community after being absent. Additionally, the hyperparameter for the Dirichlet distribution given the real dataset can be given as an input.

The ‘prop_change’ input can be used to directly incorporate longitudinal trends into the count data. The input is a vector of length $T - 1$ and is used to indicate the weights described in section 2.2.2 for the simulation framework. As described in section 2, only a subset of OTUs are modified by these weights. Three input parameters in the SimOTUCounts function are used to create the subset of modified OTUs $S$. The first input, ‘lambda’, is the threshold used to separate common and rare OTUs (default is 0.01), while the second input ‘pert_group’ determines if the common, or rare, OTUs should belong in $S$ (default is common). Finally, the proportion of the selected group of OTUs to belong to $S$ is determined by ‘eta’ (default is 0.5). In practice, the default input values to create $S$ in the function constructs a set that is half of the OTUs that have a relative abundance greater than 0.01 (common OTUs).

Additional methods exist to incorporate longitudinal trends in the simulated count matrix. Longitudinal trends can be introduced indirectly through (1) the average number of absent species at each timepoint and (2) the probability an OTU re-enters the microbiome after being previously absent. The funciton input 'zero_adjust' controls longitudinal trends in zero counts through a vector of length $t$, where each element corresponds to $o_t$, as defined in section 2.4.1. The 'zero_adjust' parameter allows a direct way to force increased, or decreased, alpha diversity over time. Alternatively, the probability of an OTU re-entering the microbiome differs by common/rare classification, as mentioned in section 2.4.2. The inputs 'non_zero_rare_prob' and 'non_zero_common_prob' represent the probability of success in a Bernoulli distribution when deciding if a rare, or common, OTU should re-enter the microbiome. If these values are relatively small (e.g. less than 0.2), there is a stronger autocorrelation structure within an individual.

SimOTUCounts() returns a list of 5 objects. The first object is the simulated count matrix with columns for individual, timepoint, and each OTU. The rows of the matrix represent each observational unit, thus the dimension of the count matrix are $nT \times (M + 2)$. The second object is a matrix of compositional vectors $\boldsymbol{a}_{it}$ (dimensions $nT \times M$). The next three objects are lists of OTUs providing information about: (1) which OTUs were perturbed, (2) which OTUs were classified as having common relative abundance, and (3) which OTUs were classified has having rare relative abundance.

### S.2.2 Visualizing longitudinal microbiome data

Several functions in the package construct visualizations to better assess real or simulated datasets. Stacked bar charts can be generated using the stacked_bar() function to visualize the average relative abundance of each OTU across time. OTU counts are also used to construct binned boxplots using the props_boxplot() function. This plot places OTUs within an individual to different classes, or bins, based upon the relative abundance of the OTU to help visualize the amount of rare, common, or absent OTUs in the dataset. Two functions are provided to construct heatmaps representing the dissimilarity between pairs of microbiomes using beta diversity values. The beta_div_hmap() function creates a heatmap comparing the beta diversity values for all pairs of observations within a subset of individuals. Alternatively, the avg_wit_beta_div_hmap() function creates the average beta diversity heatmap for all observations within an individual. Both beta diversity plots reveal longitudinal trends, such as autocorrelation, that occurs between microbiomes. The former function also allows for between-individual comparisons in microbiome samples.
